# Using Conspecific Playbacks to Investigate Contact Calling in Southeast Alaskan Humpback Whales

**DOI:** 10.1101/2021.08.25.457675

**Authors:** Michelle E.H. Fournet, Leanna P. Matthews, Annie Bartlett, Natalie Mastick, Fred Sharpe, Laurance Doyle, Brenda McCowan, James P. Crutchfield, David K. Mellinger

## Abstract

Humpback whales (*Megaptera novaeangliae*) produce calls across age and sex class and throughout their migratory range. Despite growing interest in calling behavior, the function of most calls is unknown. Among identified call types, the ‘whup’ is ubiquitous, and innate, and may serve as a contact call. We conducted an acoustic playback experiment combined with passive acoustic monitoring and visual observations to test the function of the whup on a Southeast Alaskan foraging ground. Using a before-during-after design, we broadcasted either a control sound or a unique whup call sequence. We investigated the change in whup rates (whups/whale/10 minutes) in response to treatment (whup or control) and period (*before*, *during*, or *after*). In 100% of the conspecific trials, whup rates increased during broadcasts, and whup rates were significantly higher than in *before* or *after* periods. There was no significant difference in whup rates between before and after periods during conspecific trials. In control trials, there were no significant differences in whup rates between *before*, *during*, or *after* periods. Neither whups nor control playbacks elicited an approach response. Humpback whale vocal responses to whup playbacks suggest that whups function as a contact call, but not necessarily as an aggregation signal.

## Introduction

Contact calls – acoustic signals used to make or maintain contact with conspecifics – are widespread among social vertebrates (*e.g.,* bats^12^, baboons^3^, birds^4,5^, elephants^6^, cetaceans^7^). This is particularly true in environments where visibility is limited, but information about conspecific identity or location is valuable. For marine animals that inhabit opaque waters and migrate vast distances, sound is the only viable means for detecting conspecifics beyond a few meters range^8,9^. Complex sociality has evolved in many marine mammal species^10^, and, in these groups, acoustic communication including contact calling facilitates social relationships^7^.

Humpback whales (*Megaptera novaeangliae)* are highly vocal baleen whale species with a fission-fusion social structure^11–13^. At high latitudes, humpback whales typically forage alone or in groups of 2-3 individuals^14,15^. Select individuals, however, convene in coordinated foraging groups of as many as 20 or more individuals^16,17^; membership in these large groups is stable over time^17^. Long-term social bonds between individuals of both sexes have been observed on foraging grounds, with some relationships also persisting on breeding grounds^18–21^. Strong evidence of social niche partitioning has emerged humpback whales in the Northern British Columbia - Southeast Alaskan Stock. Notably, intra-group bonds can last for over a decade^21^.

The humpback whale ‘whup’ call, is a low-frequency amplitude-modulated call with a terminal upsweep (Figure 1). Whups are produced by humpback whale populations globally, persist across generational time, and are thought to be innate in this species^22–26^. Based on use patterns, including counter calling in Southeast Alaska, Wild and Gabriele proposed that whups serve a contact function^27^. For humpback whales on foraging grounds, contact calling may be an essential tool for identifying conspecifics. It may facilitate joining events, adjust spacing to avoid resource competition, or some combination of the two based on social and ecological conditions.

**Figure 1.**
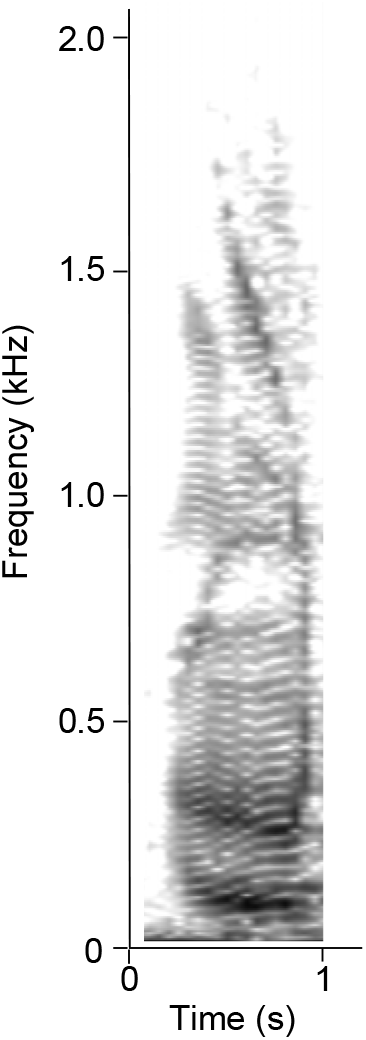
Humpback whale whup call recorded. Hann window, 2514 samples, DFT 8192, with a resolution of +-2.93 Hz, 50% overlap.

Acoustic playbacks are an effective method for investigating acoustic communication, including contact calling in marine mammals (see Deecke et al. 2006 for review^28^). To guide experimentally determining call function, robust hypothesis testing requires (1) a good description of a species’ call repertoire and social behavior, (2) an understanding of a species’ baseline acoustic behavior at sufficient detail to detect quantifiable change, and (3) sufficient high-quality recording replicates and effective knowledge for biologically-plausible call broadcasting^28^. In keeping with these recommendations, the Southeast Alaskan humpback whale call repertoire is well described, with the whup being among the most well described call types globally^24,26,27,29–31^. Because of the historic research done in Southeast Alaska, sufficient independent replicates of whup calls were available for playbacks. As a result, biologically plausible playback patterns were developed that emulate whup use in wild whale.

Despite decades of research, our understanding of humpback whales call function, and cetacean call function more broadly, remains poor. Interpreting a species’ life history requires understanding the fundamental elements of how that species interacts with the world. We offer the following study as a step toward meeting this goal.

## Methods

### Experimental Protocol

Playback trials took place at the confluence of Frederick Sound and Stephens Passage, Southeast Alaska in August and September 2019. Trials were conducted from a 16’ inflatable vessel with a single 20 hp outboard engine. For each trial, whales were visually sighted and approached at minimal speed. Ventral flukes were photographed for individual identification. We attempted to photograph every whale within an aggregation. However, for large (>10) groups of quickly moving whales, at least half of the individuals, or no less than eight individuals for very large groups, were photographed. Photographs were reviewed daily and a study catalogue was developed for rapid in field identification. Every effort was made to include whales in a unique, single playback trial for the entirety of the study. If an individual whale was identified that had been previously subjected to playback stimulus, the trial was terminated and new focal animals were pursued.

Following photo identification, a four-element drifting hydrophone array was deployed in an approximately 1-square km configuration around a focal aggregation. The research vessel was then positioned in the center of the array with the engine off and the animals were given 20 minutes to acclimate. A 10-minute *before* period followed the 20-minute acclimation. During the *before* period, we recorded bearing, behavior, and group size for each pod in visual range (to the horizon’s edge, ~5 km maximum visual distance) using a TruPulse360 (Laser Technology Inc) laser rangefinder with an internal compass. Distance to each pod was either measured with the range finder or estimated by trained observers and aggregated into one of the following bins: 0-100 m, 100-500 m, 500-1,000 m, 1,000-2,000 m, or 2,000-5,000 m. Observers were tested daily on distance accuracy by estimating distance to a kayak at randomly spaced intervals between 50 and 2000 m. Distances to the kayak were corroborated with the rangefinder. Observers accurately placed the target into the correct distance bin greater than 95% of the time.

The *before* period was followed by a 10-minute *during* period, in which a playback stimulus was projected from a EV UW30 underwater speaker (Lubell Labs) with source levels of between 136 and 142 dB_RMS_ re 1 μPa @ 1m (recording dependent). All sound files filtered in Adobe Audition based on the known broadcast parameters of the speaker to ensure that sounds were not artificially impacted by artificial amplitude modulation (See Supplementary Materials). ‘Control’ stimuli consisted of 10-minute recordings of the sounds of ambient Alaskan soundscape. ‘Whup’ stimuli consisted of one of 15 10-minute files, each containing 10 repeated whup calls at randomly spaced intervals ranging from 10 – 70 seconds. These intervals were selected based on previous analyses of *in situ* calling behavior collected in Southeast Alaska^32,33^. Whups that were included as playback stimuli were recorded in Frederick Sound, Southeast Alaska in summertime months. Recordings were unique and were not repeated during the study. Recordings were randomly selected, and observers were blind to which stimulus was played. Throughout the 10-minute during period, observers recorded the bearing, behavior, group size, and distance bin for each group of whales. The *during* period was followed by a 10-minute *afte*’ period, which was identical to the *before* period.

#### Recording Specifications

The drifting hydrophone array consisted of four independent Soundtrap ST300 hydrophones (Ocean Instruments) suspended from a surface buoy to 30 m depth. Hydrophones recorded continuously with a 24 kHz sampling rate, 16-bit resolution, and flat frequency response of ~-176 dB (instrument specific, range 174.2-176.2 dB). Hydrophones were retroactively time aligned using custom Matlab code^34^.

#### Data Processing

Whup calls were annotated by two observers (MF and AB) in RavenPro 1.6 (‘Raven’)^35^. Selections were corroborated to ensure classification consistency. Broadcasted calls were separately annotated and cross referenced to broadcast times to ensure that they were not included in analyses. Received levels for all calls were calculated using the filtered RMS amplitude measurement in Raven. Based on *in situ* received levels and reported source levels for humpback whales in Southeast Alaska^36^, an acoustic detection range was calculated for each call. Calls that were likely to have been produced beyond the 5 km visual range of the observers were omitted from analyses.

The number of whales seen in each distance bin was aggregated by trial and period (*before*, *during*, or *after*). A total whup rate was calculated for each period as the number of whups per whale per 10 minutes. For every period, whale density in each distance bin was calculated as the number of whales per kilometer^2^.

### Statistical Analysis

A repeated measures ANOVA with a random effect of trial was used to test for the effect of treatment (whup vs. control), period (before, during, after), and an interaction term on whup rates. There was one observed outlier, but analyses run with and without this observation were biologically equivalent, and the outlier was not removed in an effort to maintain as large a representative sample as possible.

A repeated measures ANOVA with a random effect of trial number was used to test for the effect of treatment, period, and an interaction term on whale density in each of the distance bins. For all models, assumptions of normality, linearity, sphericity, and homoscedasticity were visually assessed in R 4.0.4^37^ and models were deemed appropriate. Post-hoc tests were conducted with a Bonferroni correction for multiple comparisons using the emmeans package in R^38^.

## Results

Fifteen playback trials – 7 control trials and 8 whup trials – containing 856 whup calls were included in the study. Whup rates ranged from 0 – 3.25 calls whale^−1^ 10-min^−1^ (Figure 1).

The average number of whales present within a 5 km range was 17.1 (range: 3-53); average group size was 1.6 (range: 1-9). Model results indicated that period and an interaction between period and treatment significantly impacted whup rates (period: F_2,26_= 4.1. p=0.03; treatment*period: F_2,26_=5.1, p=0.01). Post-hoc tests revealed that in whup trials, whup rates were significantly higher in the *during* period than in the *before* (t_26_=3.92, p=0.0086) and *after* (t_26_=3.4, p=0.0315) periods; there were no significant differences between whup rates in before and after periods. In 100% of the whup trials, whup rates were highest in the during period. There were no significant differences in whup rates between periods in the control treatment (p=1.0) (Figure 2). There were no significant relationships between whale density and either treatment or period for any distance bins (p>0.06, supplementary material).

**Figure 2.**
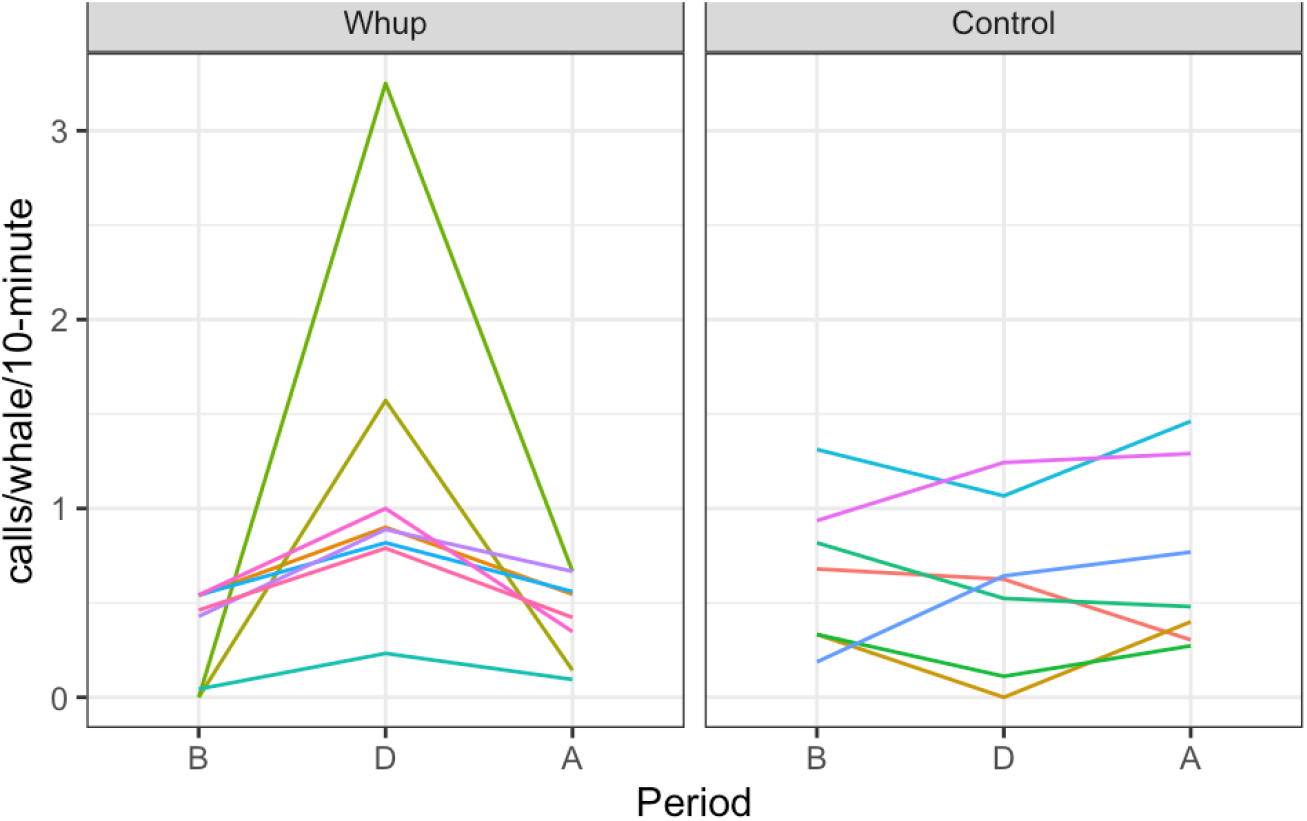
Whup rate (call/whale/10-minute) by trial period (before, during, after) and treatment (control, whup). Each uniquely colored line represents a single three-part trial.

## Discussion

In this study, we demonstrate that projecting a whup call resulted in an acoustic call-back response but did not necessarily result in an approach (which would have been indicated by an increase in whale density nearer to the playback speaker). Despite the small sample size, the acoustic response to conspecific playbacks was absolute and obvious. This is consistent with the hypothesis that the whup call is a contact call in the context of humpback whale foraging and social behavior.

Humpback whales on foraging grounds typically forage alone or in small groups^12^. In prey rich habitats like Southeast Alaska, excessive crowding of a resource is at odds with the competitive assumptions of optimal foraging theory, which predicts that animals adopt strategies that maximize prey consumption while minimizing energy expenditure^39,40^. By this reasoning, when prey are abundant, contact calls should function to space individuals, avoiding unnecessary energetic expenditures associated with resource competition. Prey were abundant during this study; krill or herring were noted daily. Thus, the sound of a calling conspecific in this ecological context may be unlikely to elicit an approach response lest crowding ensure. This would be particularly pertinent if the signaler and the receiver lacked a strong social affiliation that may otherwise lead to group foraging.

In Southeast Alaska, bubble-net foraging is characterized by individuals or groups deploying bubble-nets in tandem with a highly stereotyped vocalization to forage on Pacific herring (*Clupea pallasii*)^16,41,42^.

Bubble-netting is a known driver of large group formation and social relationships ^17,21^; whales that participate in bubble-netting exhibit stronger social preferences, longer social bonds, and greater social selectivity than those who do not^17,21^. In this study, the social relationship between the broadcast whale and the receiving whale – including if one or both had a previous relationship of bubble-netting together – may have influenced the probability of joining. Whales with coordinated foraging history may maximize energetic intake by joining another group forager, while non-group foragers may avoid individuals with whom they lack a social relationship to avoid resource competition. It is unknown whether the recordings used in this study came from individuals who engage in bubble-net feeding. That said, for the decision to join or not to join to occur, contact must first be made between individuals.

The observed increase in whup rates associated with conspecific playbacks in this study is evidence that whup calls facilitate contact between individuals. Given this, it is probable that whup calls mediate social relationships in humpback whales. For long term bonds to persist, there must be some cue to facilitate individual recognition. Across taxa, contact calls contain vocal signatures that receivers use to make decisions. Spider monkeys (*Ateles geoffroyi*) use contact calls to achieve flexibility in spacing while maintaining specific social relationships^43^. Pallid bats (*Antrozous pallidus)* reply to and approach known individuals, but are unlikely to approach unknown conspecifics^44^. Similarly, contact calls of North Atlantic right whales (*Eubalaena glacialis*) serve a long range contact function^45,46^ and are individually distinct^47^. Our study supports the view that individual identity is acoustically encoded in whup calls. This was qualitatively observed during analysis and is the subject of future research.

Notably, our results differ from results of a humpback whale playback study conducted in Hawaii in 1985-1986^48^. The Hawaiian experiment, which did not include the whup, found that the most pronounced response was a rapid approach to the playback of Alaskan feeding calls. This ‘charging’ behavior was notably absent in our study, indicating a strong functional difference between whup calls and feeding calls. The function of feeding calls is now well described^41,42,49^, but the differential response to the playback of whups versus feeding calls is evidence that humpback whales can and do, differentiate between different social calls.

This study corroborates observed calling patterns in Southeast Alaskan humpback whales^27^ and confirms the hypothesis that whups serve a contact calling function. Dedicated studies into cetacean communication are valuable for both understanding basic ecology and behavior as well as framing management and conservation action in light of a species’ life history. Understanding the impacts of anthropogenic noise and other human disturbances on humpback whales and other marine mammals is contingent on understanding the role that sound plays in the lives of these species. This work supports that broader goal.

## Acknowledgments

The authors thank Debbie Koyler of the SETI Institute, Don Holmes for field support, Holger Klinck for array design support, and Dean Dotson for facilities use. Thanks to Danielle Nelson for coding assistance, D. Brixey for logistical support, and J.F. Criminey for analysis assistance. This project was supported by a grant to the SETI Institute from the Templeton World Charity Foundation, Inc (Laurance Doyle, PI). The opinions expressed in this publication do not necessarily reflect the views of Templeton World Charity Foundation, Inc. This is PMEL contribution number XXXXXX. Research was conducted under NMFS Permit #19703.

## Supplemental Material

**Table 1.**
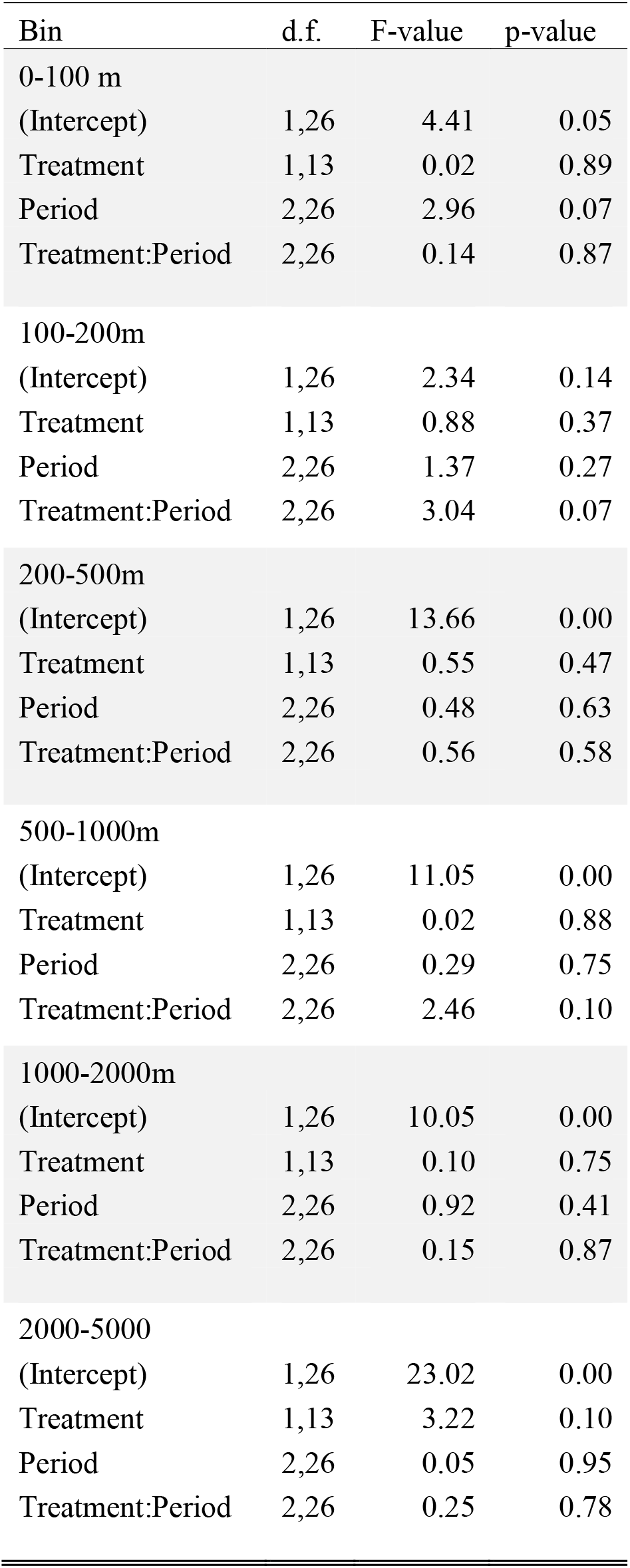
Model results for repeated measure ANOVA with a random effect of trial testing for the effect of period (before, during, after), treatment (control, whup), and an interaction on whale density. Bin indicates which distance bin the model refers to.

### Filtering Technique

All sound files originated with a flat frequency response based on hydrophone frequency response curves at the relevant bandwidth (50-500 Hz). An inverse filter was manually created and applied (Figure 1) using Adobe Audition to account for the frequency response of the UW20 underwater speaker. The inverse filter only accounts for the speaker frequency response up to 1 kHz, as all of the recordings used for this study were lower than 500 kHz.

**Figure 1.**
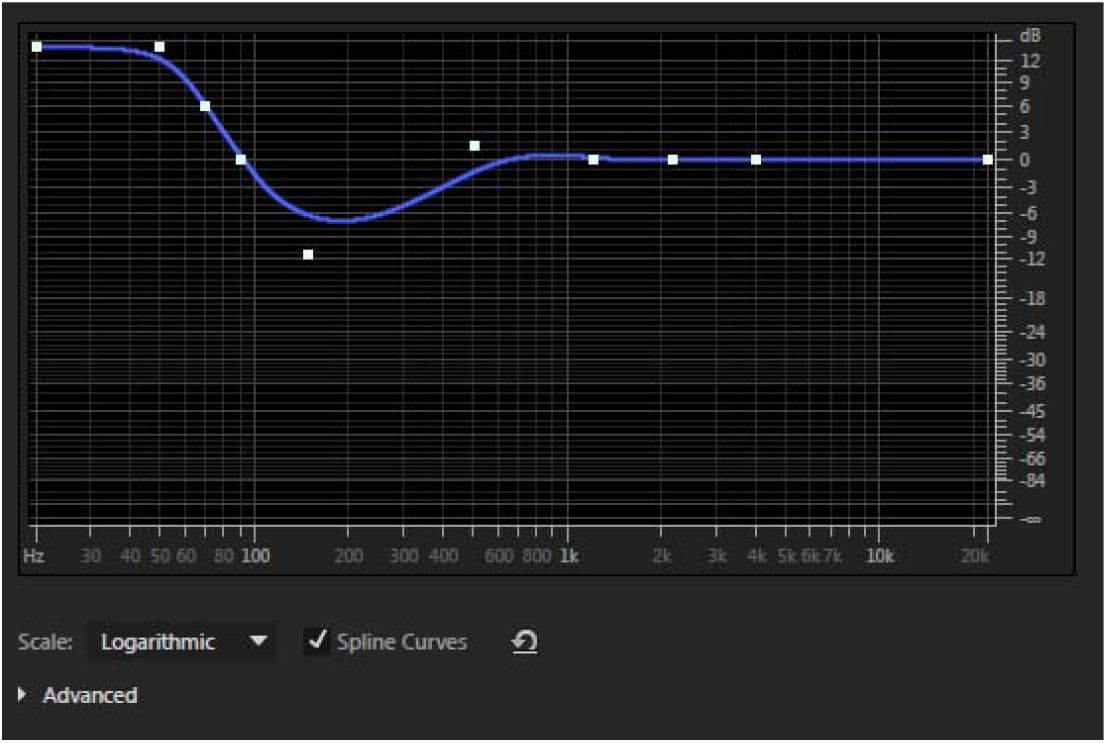
Inverse filter applied to playback sound files using Adobe Audition to result in flat playback amplification.

